# Membrane permeabilization is mediated by distinct epitopes in mouse and human orthologs of the necroptosis effector, MLKL

**DOI:** 10.1101/2021.05.03.442385

**Authors:** Ashish Sethi, Christopher R. Horne, Cheree Fitzgibbon, Karyn Wilde, Katherine A. Davies, Sarah E. Garnish, Annette V. Jacobsen, André L. Samson, Joanne M. Hildebrand, Ahmad Wardak, Peter E. Czabotar, Emma J. Petrie, Paul R. Gooley, James M. Murphy

## Abstract

Necroptosis is a lytic programmed cell death pathway with origins in innate immunity that is frequently dysregulated in inflammatory diseases. The terminal effector of the pathway, MLKL, is licensed to kill following phosphorylation of its pseudokinase domain by the upstream regulator, RIPK3 kinase. Phosphorylation provokes the unleashing of MLKL’s N-terminal four-helix bundle (4HB or HeLo) domain, which binds and permeabilizes the plasma membrane to cause cell death. The precise mechanism by which the 4HB domain permeabilizes membranes, and how the mechanism differs between species, remains unclear. Here, we identify the membrane binding epitope of mouse MLKL using NMR spectroscopy. Using liposome permeabilization and cell death assays, we validate K69 in the α3 helix, W108 in the α4 helix, and R137/Q138 in the first brace helix as crucial residues for necroptotic signaling. This epitope differs from the phospholipid binding site reported for human MLKL, which comprises basic residues primarily located in the α1 and α2 helices. In further contrast to human and plant MLKL orthologs, in which the α3-α4 loop forms a helix, this loop is unstructured in mouse MLKL in solution. Together, these findings illustrate the versatility of the 4HB domain fold, whose lytic function can be mediated by distinct epitopes in different orthologs.

## INTRODUCTION

Necroptosis is a caspase-independent, lytic cell death mode with ancestral origins in host defense^1, 2, 3, 4, 5, 6, 7, 8, 9^, which is frequently dysregulated in disease^10, 11, 12, 13, 14, 15, 16^. The inflammatory nature of necroptosis has led to its implication in a range of human pathologies, including renal^11, 13^ and gastrointestinal diseases^14, 17^. Necroptotic signaling is instigated by ligation of death receptors, such as TNF Receptor 1, or pathogen detectors, such as ZBP1/DAI, in cellular contexts where the IAP E3 Ubiquitin ligase family and the proteolytic enzyme, Caspase-8, are downregulated or their catalytic activities compromised (reviewed in ref.^18^). Downstream of receptor activation, a cytoplasmic platform termed the necrosome is assembled in which the Receptor-interacting serine/threonine protein kinase (RIPK)-1 recruits RIPK3, leading to its activation by autophosphorylation^1, 19, 20, 21, 22^. Subsequently, RIPK3 phosphorylates the pseudokinase domain of the necroptotic executioner, Mixed lineage kinase domain-like (MLKL)^23, 24^, to induce its dissociation from the necrosome^25, 26, 27^, assembly into high molecular weight complexes^27, 28, 29^, and trafficking to the plasma membrane^18, 27, 30^. When a threshold level of activated MLKL accumulates at the plasma membrane^27, 30^, MLKL perturbs the lipid bilayer to cause cell death via an incompletely understood mechanism^31^. This mode of cell death involves the leakage of cellular contents, including DAMPs, into the extracellular milieu to provoke an inflammatory response^32^.

While the core principles of necroptotic signaling and MLKL activation are preserved between species, the precise molecular mechanisms appear to differ. Detailed studies of the pseudokinase domains of MLKL orthologs have revealed their propensity to adopt distinct conformations^23, 29, 33, 34, 35^, which governs recognition by RIPK3 and results in very strict species specificity^36, 37^. Additionally, the role of activation loop phosphorylation in triggering MLKL activation appears to vary between orthologs. Phosphorylation serves as a trigger for release of the killer 4HB domain in mouse MLKL^23, 28, 38, 39^, as a cue for MLKL release from the necrosome and interconversion to the closed, active form in human MLKL^25, 26, 29, 34^, and a likely role in negating occupation of the pseudoactive site in horse MLKL^33^. Consistent with the diverse regulatory mechanisms governing the MLKL pseudokinase domain molecular switch, the N-terminal executioner 4HB domain exhibits heterogeneous membrane permeabilization between species^37^. However, the molecular basis for how and why recombinant mouse MLKL 4HB domain more efficiently permeabilizes lipid bilayers than the human and chicken 4HB domains has remained unclear.

NMR studies of human MLKL’s 4HB domain over the past 7 years have provided important insights into the residues involved in phospholipid headgroup and inositol phosphate recognition^40, 41, 42, 43^. These studies have implicated basic residues located principally within the α1 and α2 helices in negatively-charged phospholipid binding^42, 43^, while inositol phosphate recognition relies on an epitope centred on the loop connecting the α2 and α3 helices, and the α1 helix, including lipid binding residues^40, 41^. Considering the low sequence identity between human and mouse MLKL 4HB domains^33, 44^, we employed NMR spectroscopy to define the residues that mediate lipid recognition in mouse MLKL and to identify structural differences from its human counterpart. Remarkably, in NMR relaxation experiments, we identified residues on the opposing face of the mouse MLKL 4HB domain, relative to those implicated in human MLKL lipid and inositol phosphate recognition, as mediators of liposome binding. Mutation of these residues compromised liposome permeabilization *in vitro*, with a subset of these sites found to be functionally crucial for mouse MLKL necroptotic signaling in cells. Collectively, these data illustrate that mouse and human MLKL rely on distinct lipid-binding residues to enact cell death and support the idea that the 4HB (also known as HeLo) domain can serve as a versatile scaffold for lipid recognition and permeabilization.

## RESULTS

### Mouse MLKL 4HB domain adopts a folded helical structure

To characterize the structure of N-terminal four-helix bundle (4HB) domain and the first brace helix of mouse MLKL (residues 1-158; termed mouse MLKL_(1-158)_ herein) in solution, we subjected ^2^H,^15^N,^13^C-labelled protein to non-uniformly sampled three-dimensional NMR methodology. As previously, mouse MLKL_(1-158)_ purified as a monomer, owing to the absence of the second brace helix that is required for trimerization ^36^. From these 3D NMR experiments, we could successfully assign 95% of the backbone resonances corresponding to residues 1-158 of mouse MLKL (**Figure 1a**), with additional backbone amides corresponding to the vector-encoded remnant sequence (GAMGS) also observed (numbered as residues −4 to 0 in **Figure 1b-c**). As anticipated from the crystal structure of full-length mouse MLKL, the 4HB domain (residues 1-125) and the adjacent brace helix 1 exhibited predominantly positive ΔCα−ΔCβ smoothed values (**Figure 1b**), which is consistent with the expected helical structure. In contrast, the region corresponding to the S82 to G91 backbone amides exhibited trends to both negative and positive ΔCα−ΔCβ values (ΔCα−ΔCβ < ±1.0), indicating a lack of regular secondary structure in this region. Interestingly, the overlapping region encompassing S79 to K94 could not be modelled in the mouse MLKL crystal structure due to lack of electron density^23, 45^, consistent with it occurring as an unstructured loop. To further investigate the internal dynamics within the mouse MLKL 4HB domain structure in solution, we recorded a steady state ^15^N{^1^H}NOE experiment^46^ on ^15^N-labelled mouse MLKL_(1-158)_ (**Figure 1b**). The average ^15^N{^1^H}-NOE values (0.82±0.07) for mouse MLKL (residues D2-V150) support the existence of the mouse MLKL 4HB domain and adjacent brace helix occurring in solution as a structured protein with flexible termini (vector encoded residues −4 to 0 and residues 151-158, average ^15^N{^1^H}-NOE values < 0.5). For the region S79 to K94, which corresponds to the loop connecting the α3 and α4 helices in the mouse MLKL crystal structure, an average ^15^N{^1^H}-NOE value of (0.62±0.05) was recorded. These data are consistent with this loop exhibiting higher flexibility than the remaining mouse MLKL 4HB+brace core structure.

**Figure 1.**
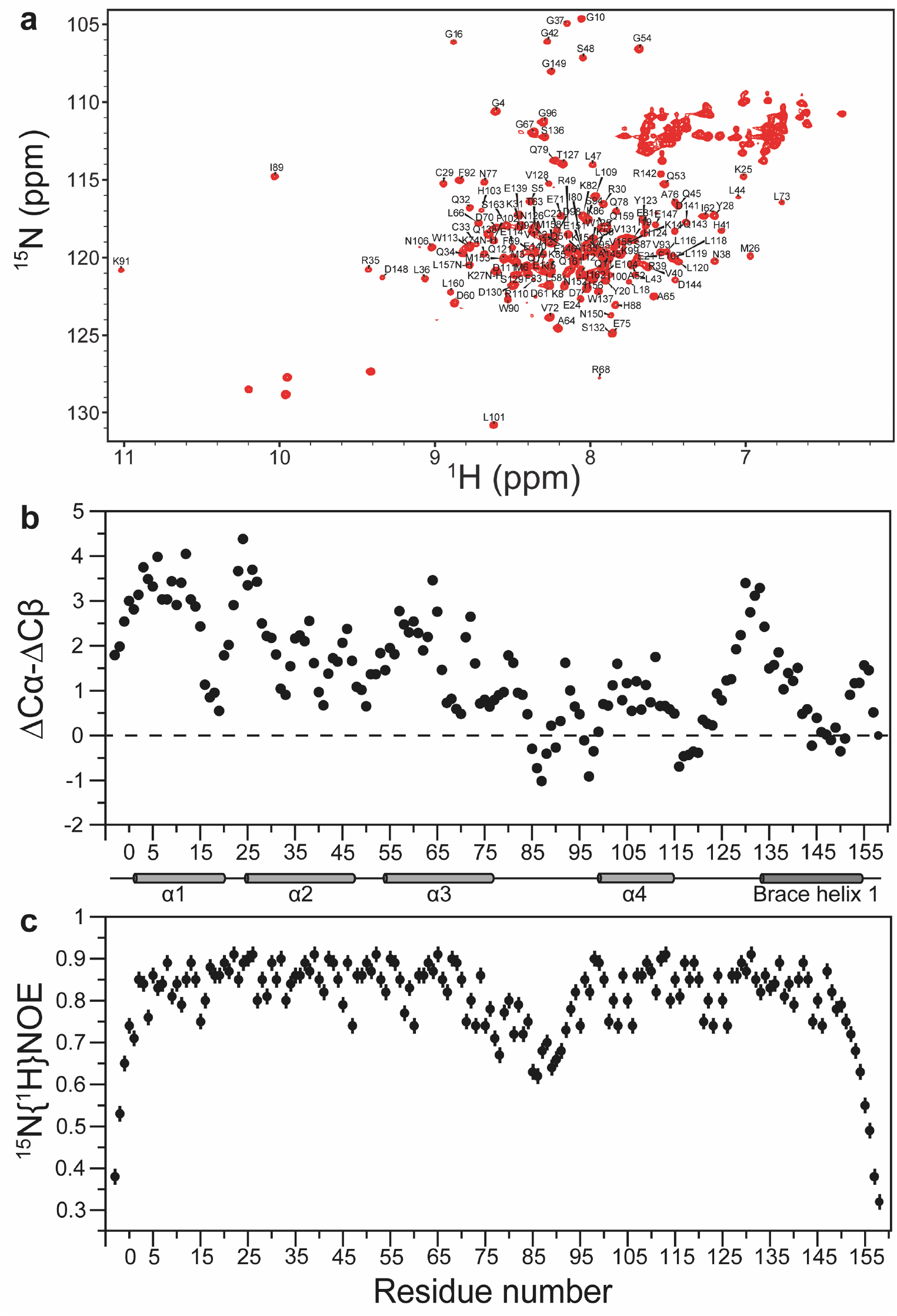
Mouse MLKL adopts a folded helical structure. **a)** Full ^1^H, ^15^N HSQC spectrum of mouse MLKL_(1-158)_ with residue assignments. Unmarked resonances in the upper right corner belong to Asn and Gln sidechains and in the lower left corner belong to Trp-indole sidechains have not been assigned. The residues, GAMGS, are a cloning artifact and numbered −4 to 0, while residues M1-S158 are from mouse MLKL. **b)** Plots of ^13^Cαβ secondary chemical shifts and **c)** Steady state ^15^N{^1^H}-NOEs for mouse MLKL_(1-158)_ with error bars calculated based on average estimated noise level for ^15^N{^1^H}-NOE. Experiments were conducted at pH 6.8 and 25 °C. The helical secondary structure shown between panels (**b**) and (**c**) reflects that from the full-length mouse MLKL crystal structure (PDB, 4BTF)^23^.

### Two clusters mediate mouse MLKL 4HB+brace binding to liposomes

Studies of the human MLKL 4HB domain by NMR spectroscopy have identified principally basic residues as the mediators of lipid binding, lipid permeabilization and cell death^42, 43^. While analogous studies have not been performed to date on mouse MLKL 4HB domain, very few of the key residues within human MLKL are conserved in the mouse ortholog. Accordingly, we sought to deduce which mouse MLKL 4HB domain residues mediate lipid binding, and whether they spatially differ to the reported lipid-binding residues in the human MLKL 4HB domain, by performing a 2D ^1^H-^15^N HSQC monitored titration of uniformly ^15^N-labeled mouse MLKL_(1-158)_ with liposomes of a plasma membrane-like composition. Using this approach, we identified two clusters of residues in mouse MLKL that exhibited marked attenuation of peak intensity (**Figure 2a-b**). Diminished peak intensity is a sensitive means of detecting the engagement of different sites within the mouse MLKL 4HB+brace protein with liposomes, which enables each individual backbone amide resonance to serve as a probe to report changes in their solvent exposure, motions and interactions. Among these two clusters, cluster I comprised R34-Q40 in the α2 helix and D106-E110 in the neighboring α4 helix; and cluster II was composed of V67-A71 in the α3 helix, N92-N101 in the α4 helix and preceding region, and D136-D139 in brace helix 1 (**Figure 2b**). We further validated clusters I and II as liposome interacting sites in mouse MLKL using a ^15^N amide spin transverse relaxation (^15^N-R_2_) experiment at 70.9 MHz for mouse MLKL_(1-158)_ in the presence and absence of liposome in the ratio 1:0.5 (mouse MLKL:liposome) at 25 °C (**Figure 2c**). As expected, in the presence of liposomes, the ^15^N-R_2_ values for regions clusters I and II within mouse MLKL_(1-158)_ (**Figure 2c**) showed a marked increase, reflecting the chemical exchange on a fast timescale with liposomes. Collectively, these data confirm roles for sites clustered on the centre of α2 and α4 helices (cluster I) and the N-terminal ends of the α4 helix and the flanking α3 and brace helices (cluster II) in liposome binding, suggesting liposome engagement is mediated via an extended interface (**Figure 2b**).

**Figure 2.**
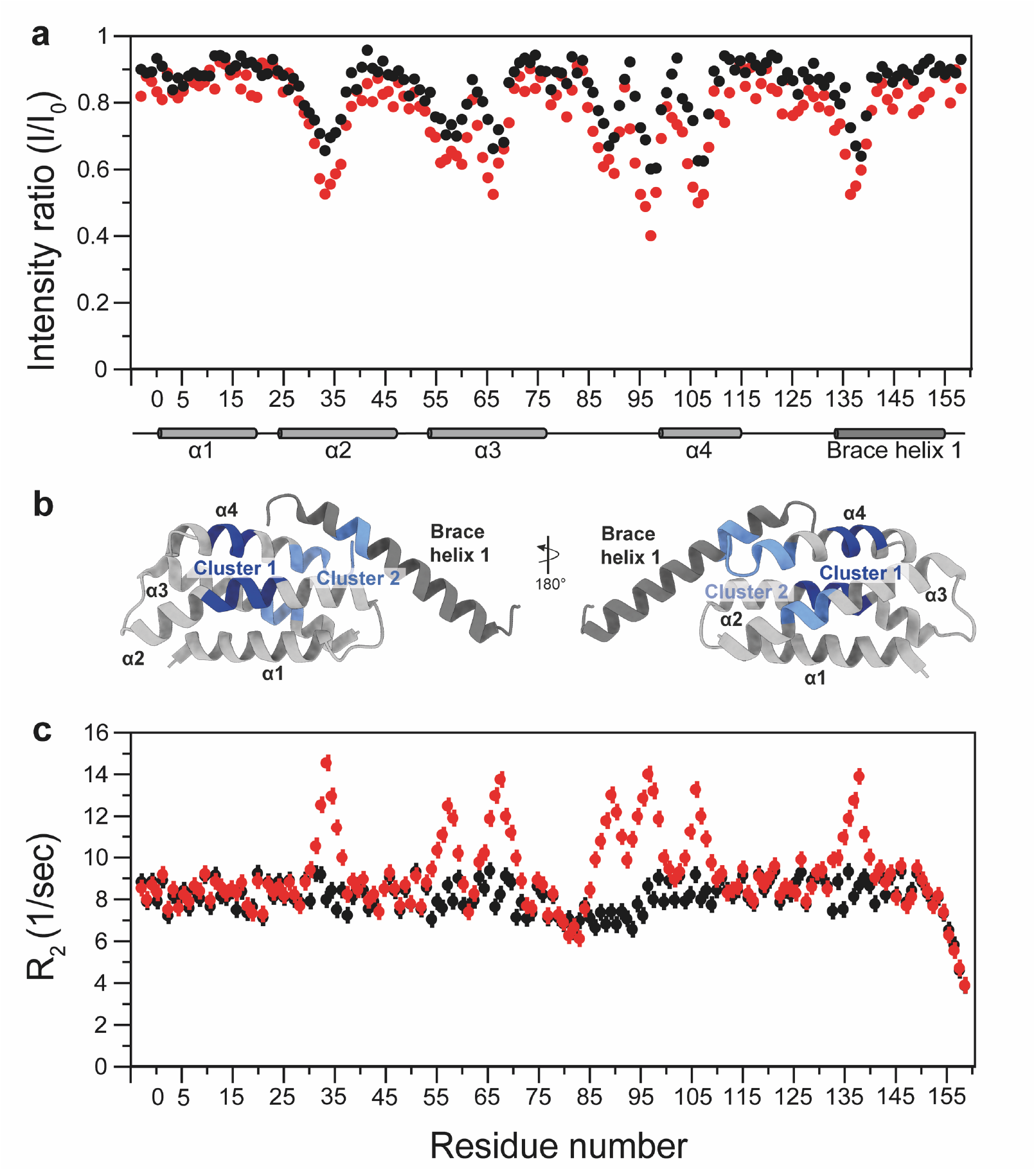
Two clusters of residues in mouse MLKL_(1-158)_ mediate lipid binding. Mouse MLKL_(1-158)_ binds to liposomes shown by titrating 100 μM ^15^N-labelled mouse MLKL_(1-158)_ with liposomes in the 1:0.5 (black circles) and 1:1 (red circles) ratio. **a)** Plot of change in ^1^HN and ^15^N peak intensity, where I is the intensity of the peak in the presence of liposome and I_0_ in the absence. The helical secondary structure shown between panels (**a**) and (**b**) reflects that from the full-length mouse MLKL crystal structure (PDB, 4BTF)^23^. **b)** Clusters I (light blue) and II (dark blue) of residues in mouse MLKL_(1-158)_ that exhibited marked attenuation of peak intensity are mapped onto the mouse MLKL crystal structure (PDB, 4BTF). **c)** Backbone ^15^N transverse relaxation rates (R_2_) measured at ^15^N frequency of 70.9 MHz, with liposomes in the 1:0.5 (black circles) and 1:1 (red circles) ratio. Error bars were calculated using Monte Carlo Simulations for R_2_ measurement. Experiments in (**a**) and (**c**) were conducted at pH 6.8 and 25 °C.

### Membrane binding residues contribute to liposome permeabilization

We next sought to examine whether individual substitutions of sites identified as liposome interactors in NMR spectroscopy experiments would impact liposome permeabilization. To this end, we introduced Ala substitutions to H36 (α2 helix), K69 (α3 helix), N92, H98, W108 (α4 helix and preceding region) and R137/Q138 (brace helix 1) (**Figure 3a**) in mouse MLKL_(1-158)_ (**Figure 3b-c**) and prepared recombinant proteins for *in vitro* dye release assays. These residues were selected because they comprise the solvent-exposed sites in clusters I and II when mapped to the mouse MLKL crystal structure^23^. We reasoned that shifts observed for adjacent hydrophobic core residues were likely a secondary effect of their proximity to lipid-binding residues and therefore did not mutate core residues to avoid disrupting 4HB domain folding. In these assays, liposomes loaded with the self-quenching dye, 5(6)-carboxyfluorescein, were incubated with 8 μM recombinant mouse MLKL_(1-158)_ and dye release measured spectrophotometrically. While wild-type mouse MLKL_(1-158)_ permeabilized liposomes with comparable kinetics to previous studies^36, 37^, alanine substitutions of sites identified as liposome binding residues in NMR experiments led to dampened permeabilization of liposomes in all cases except N92A (**Figure 3d-e**). Notably, alanine substitution of the neighbouring residues, H98 and R137/Q138, within cluster II compromised liposome permeabilization most severely. These data validate residues located on the α2, α3 and α4 helices and the first brace helix as liposome interactors, which individually likely contribute to membrane permeabilization. It is noteworthy that the introduction of individual mutations into mouse MLKL_(1-158)_ did not lead to complete abrogation of liposome permeabilization, as expected based on earlier studies of the human MLKL ortholog^43, 47^. As in human MLKL, we expect that residues on mouse MLKL_(1-158)_ act collectively to mediate lipid binding and bilayer permeabilization, and as such, there is some redundancy between lipid-interacting residues within the domain.

**Figure 3.**
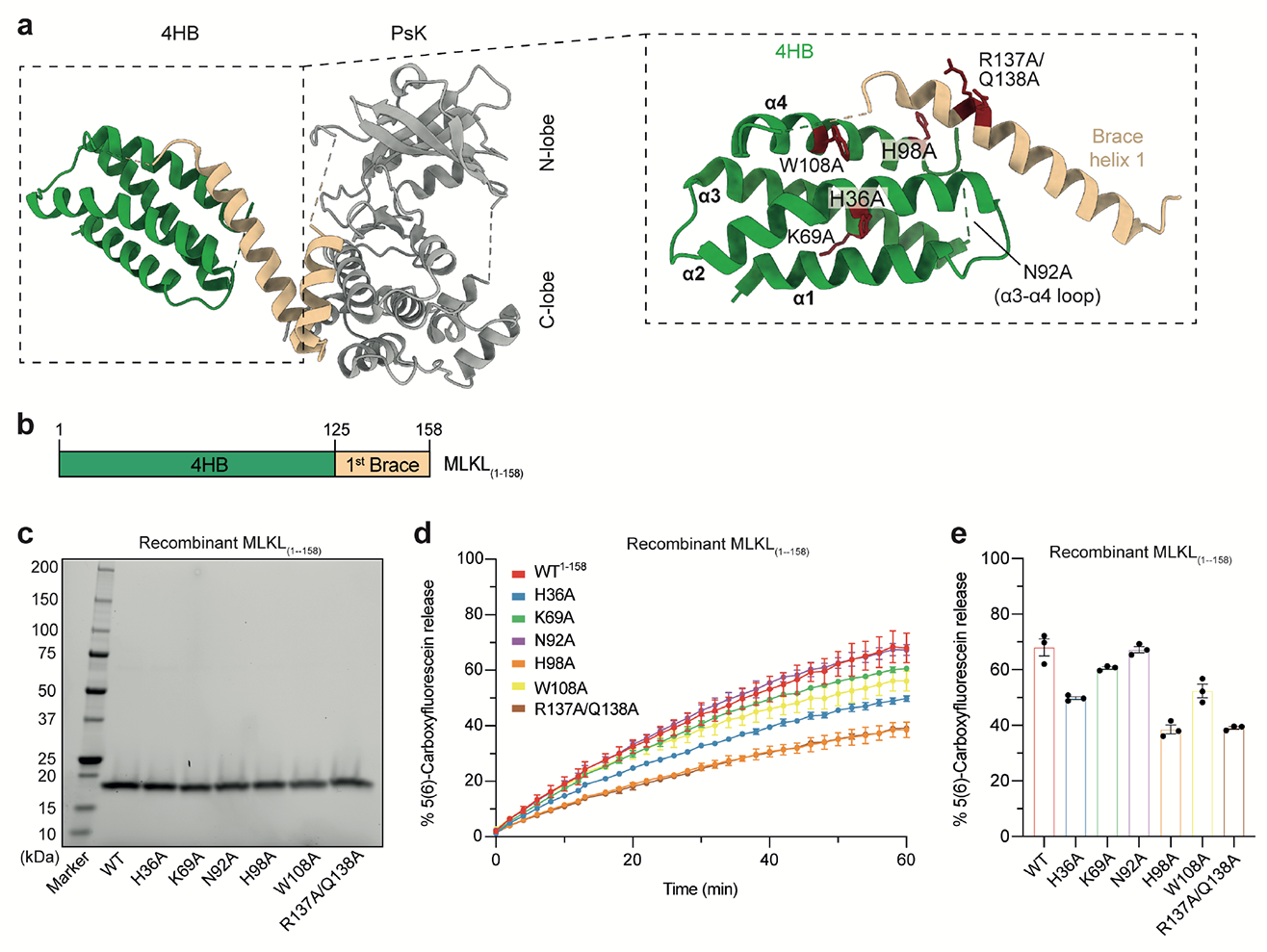
Liposome permeabilization assays validate lipid-interacting residues. **a)** Full-length mouse MLKL crystal structure (PDB, 4BTF)^23^, comprising a 4HB (green), two-brace helices (beige) and a bilobal pseudokinase (PsK) domain (grey). The surface-exposed residues implicated to interact with lipids using NMR spectroscopy are shown as sticks and coloured red (inset). Ala substitutions were introduced to each of these residues in MLKL_(1-158)._ **b**) Architecture of mouse MLKL_(1-158)_. **c)** Reducing SDS-PAGE of all mouse MLKL_(1-158)_ purified constructs. Each purified recombinant protein construct was resolved by reducing SDS-PAGE to assess purity. **d)** Liposome dye release assay using recombinant wild-type and mutant mouse MLKL_(1-158)_ at 8 μM. Release of 5(6)-Carboxyfluorescein was monitored at 485 nm over 60 min. **e)** Evaluation of total dye release from wild-type and alanine substitution mutants of mouse MLKL_(1-158)_. Data in (**d-e**) represent mean ± SEM of three independent assays.

### Residues in the α3 helix, α4 helix, and in the first brace mediate necroptotic signaling

Having identified residues that compromise liposome permeabilization by recombinant mouse MLKL_(1-158)_, we next introduced Ala substitutions of each liposome binding residue into constructs encoding full-length mouse MLKL (**Figure 4a**). We stably introduced wild-type and mutant MLKL into *Mlkl*^*-/-*^ Mouse Dermal Fibroblast (MDF) cells via a doxycycline-inducible lentiviral system. Following doxycycline treatment to induce expression (**Supp. Fig. 1**), we examined the cellular response to the necroptotic stimulus, TSI (Tumor necrosis factor (TNF, T); Smac mimetic, Compound A (S); and the pan-caspase inhibitor, emricasan/IDN-6556 (I)^25, 36, 48^) using IncuCyte live cell imaging. The capacity of each MLKL construct to reconstitute the necroptotic signaling pathway was measured by quantifying SYTOX Green uptake, as a measure of cell death, relative to the total number of cells stained by cell-permeable DNA probe, SPY620. Importantly, expression of full-length wild-type mouse MLKL successfully restored sensitivity to the necroptotic stimulus, TSI, resulting in ∼80% cell death at 5 h post-TSI stimulation (**Figure 4b**). Comparable necroptotic cell death kinetics were also observed for the mutant MLKL constructs, H36A, N92A and H98A, indicating that, individually, these residues do not impact necroptotic signaling (**Figure 4b; Supplementary Fig. 2a-c**). In contrast, alanine substitution of K69, W108 and R137/Q138 markedly reduced cell death relative to wild-type MLKL, demonstrating that substitution of these residues attenuates necroptotic signaling (**Figure 4b-c; Supplementary Fig. 2d-f**). We further validated these differences in cell death between MLKL constructs in an orthogonal assay by quantifying the release of lactate dehydrogenase (LDH) that arises from plasma membrane lysis following TSI stimulation (**Figure 4d**). Consistent with necroptotic cell death monitored by IncuCyte imaging, the LDH release values identified K69, W108 and R137/Q138 located on the α3, α4 helices and the first brace helix, respectively, as functionally crucial for mouse MLKL necroptotic signaling.

**Figure 4.**
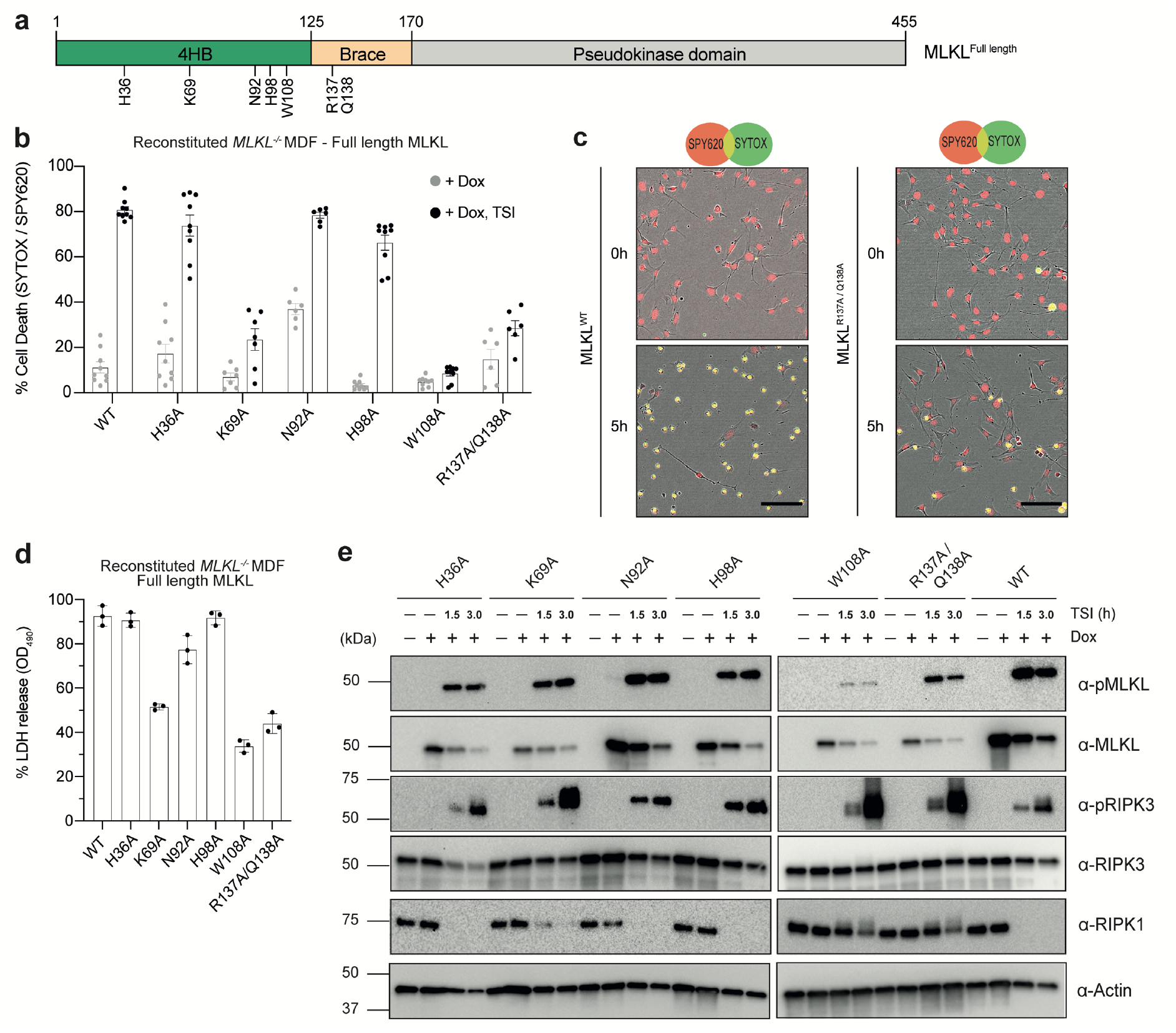
Mutation of lipid-binding residues in the 4HB and in the first brace compromise necroptotic signaling. **a)** Architecture of full-length mouse MLKL, with each Ala substitution highlighted. **b**) Evaluation of necroptotic signaling for wild-type (WT) and alanine substitution mutants of full-length mouse MLKL in MDF *Mlkl*^*-/-*^ cells. WT or mutant mouse MLKL expression was induced with doxycycline (Dox) and cell death was quantified using IncuCyte S3 live cell imaging in the presence or absence of the necroptotic stimulus, TNF and Smac-mimetic Compound A and pan-caspase inhibitor, IDN-6556, (TSI) for 5 h, by determining the number SYTOX Green-positive cells (dead cells) relative to the number of SPY620-positive cells (total cell confluency). Data represent mean ± SEM from three biologically independent MDF *Mlkl*^*-/-*^ cell lines (*n* = 6 to 9). **c**) Time-lapse micrographs of SPY620 uptake (SPY; red) and SYTOX Green uptake (SYTOX; green) as indices of total cell confluency and cell death, respectively. Both 0 and 5 h timepoints are shown post Dox induction and TSI treatment. Micrographs are representative of *n* = 6-9 independent experiments using IncuCyte S3 live cell imaging. Scale bar (black) represents 100 μm. **d)** Following induction with doxycycline and 3 h of TSI treatment, the extracellular release of lactate dehydrogenase (LDH) as an index of plasma membrane lysis (relative to detergent-treated cells) was measured by spectrophotometry at 490 nm. Data represent mean ± SD of three independent experiments. **e)** *Mlkl*^*-/-*^ MDF cells expressing WT and mutant full-length mouse MLKL following Dox induction were treated with the necroptotic stimulus, TSI, for 1.5 or 3 h. Whole cell-lysates were then fractioned by SDS-PAGE and probed by immunoblot for RIPK1, RIPK3, pRIPK3, MLKL and pMLKL with anti-actin as a loading control. Immunoblots are representative of *n* = 2 independent experiments.

MLKL binds to membranes once it has been phosphorylated by RIPK3 ^28, 39, 49, 50^. Consistent with this notion, mutation of the surface-exposed residues in clusters I and II did not have a major influence on the phosphorylation of RIPK3 or MLKL upon TSI-stimulation (**Figure 4e**). Importantly, we observed the presence of phosphorylated MLKL and RIPK3 in wild-type and all mutant MLKL constructs after TSI-treatment (1.5 or 3 h), indicating that our cellular findings are not attributable to compromise of an upstream necroptosis pathway checkpoint. We noted that while RIPK1 was detected under basal conditions for all cell lines in our immunoblots, RIPK1 was not detectable following TSI-stimulation in all conditions, which we attribute to compromised detection following the post-translational modifications that accompany necroptotic stimulation^21^, as previously reported^49^. It is notable that K69A, W108A and R137A/Q138A MLKL were expressed at lower levels than most constructs, following doxycycline treatment (**Figure 4e**). However, because these mutant MLKL proteins were expressed at an equivalent level to H36A MLKL, which exhibited comparable cell death kinetics to wild-type MLKL, any deficits in necroptotic signaling are not a consequence of lower protein expression. We used a mouse MLKL pseudokinase domain-specific antibody to detect MLKL expression (WEHI clone 5A6^49^), which ensures any differences in detection reflect levels, rather than altered reactivity that might arise from using the brace region-directed antibody (WEHI clone 3H1^23^).

## DISCUSSION

Over the past five years, it has emerged that MLKL orthologs exhibit differing propensities to permeabilize lipid bilayers and thus to enact cell death. While our understanding of the divergent activation and regulatory mechanisms among the pseudokinase domains of MLKL orthologs has been greatly enhanced by detailed structural studies^23, 29, 34, 35^, knowledge of differences between their membrane-permeabilizing executioner domain, the N-terminal four-helix bundle (4HB) domain, is limited. Distinctions between human and mouse MLKL 4HB domains are evident from their sequences, with only 52% identity at the amino acid level, and here we sought to further understand differences at the mechanistic level using a combination of NMR spectroscopy, biochemical and cellular assays.

Our NMR spectroscopy experiments validated that the mouse MLKL 4HB and first brace helix adopts a folded helical structure in solution, consistent with the fold observed in the crystal structure of full-length mouse MLKL^23^. In keeping with the full-length mouse MLKL crystal structure, we did not observe consistent positive ΔCα−ΔCβ values for residues within the loop connecting the α3 and α4 helices, indicating that this loop does not form a helix. Indeed, the reduced ^15^N{^1^H}-NOE values in this loop (relative to the domain overall) supports the assertion that the loop is flexible, as originally proposed based on the lack of density for this region in the full-length mouse MLKL crystal structure^23^. This contrasts the human MLKL 4HB domain NMR^41, 43, 51^ and crystal structures^30, 51^, where the loop connecting the α3 and α4 helices forms a helix, which is the target of the covalent MLKL inhibitor, NSA^24^. Interestingly, like human, but in contrast to mouse, MLKL 4HB domain structures, the recent cryo-EM structure of a plant MLKL ortholog, which is believed to have arisen via convergent evolution, revealed a helix in the loop connecting the α3 and α4 helices of the 4HB domain^52^.

Using NMR relaxation experiments, we then implicated two clusters of residues as lipid interactors by titrating the N-terminal helical region of mouse MLKL with liposomes that emulated a plasma membrane composition. Importantly, we chose to examine a monomeric form of the mouse MLKL (residues 1-158; 4HB domain and first brace helix)^36^ in our NMR experiments, which allows us to attribute any resonance broadening in titrations to liposome binding, and not oligomerization events. Broadly, the identified sites are spatially-proximal to those identified as key mediators of necroptotic signaling in our earlier cellular studies of mouse MLKL^28, 37^ (**Figure 5a-b; Supplementary Table 1**). However, whether the arising defects in cell signaling were attributable to deficits in lipid recognition, or other impacts on necroptotic checkpoints, had not been formally examined. Here, we add to current knowledge by establishing roles for mouse MLKL K69 (α3 helix), W108 (α4 helix) and R137/Q138 (first brace helix) in liposome permeabilization in dye release assays and necroptotic signaling in reconstituted *Mlkl*^*-/-*^ MDF cells. Our finding that, despite deficits in signaling, these MLKL constructs and the upstream regulator, RIPK3, were phosphorylated following necroptotic stimulation indicates that mouse MLKL can still undergo RIPK3-mediated phosphorylation via the proposed transient “kiss and run” mechanism^23, 26, 28, 45^. These data support the notion that the loss-of-function mutations identified here arise as a consequence of compromised lipid recognition and membrane permeabilization downstream of RIPK3 interaction.

**Figure 5.**
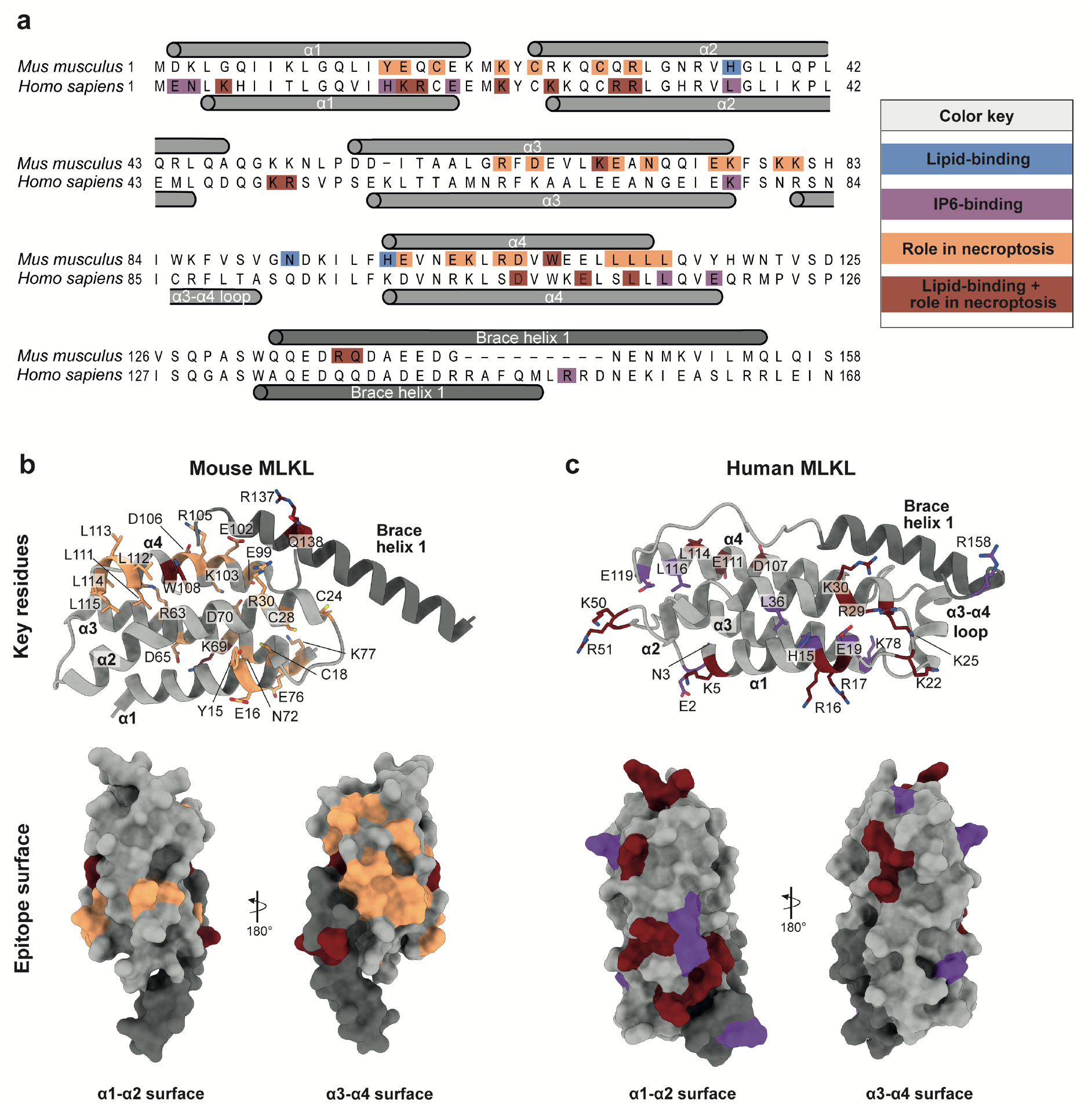
Membrane permeabilization is mediated by distinct epitopes in mouse and human MLKL. **a)** Sequence alignment of MLKL four-helix-bundle (4HB) + first brace helix from mouse and human MLKL. Secondary structure from experimental structures of mouse^23^ and human^43^ MLKL 4HB + brace helices are annotated above and below the sequences, respectively. Residues in blue have been validated to bind lipids; residues in purple have been validated to bind inositol hexaphosphate (IP6); residues in orange when mutated to alanine exhibited deficits in cellular necroptotic signaling; and mutation of lipid-binding residues in dark red exhibited deficits in cellular necroptotic signaling. **b)** Cartoon representation of mouse MLKL 4HB (grey) + first brace (dark grey, Top panel) (PDB, 4BTF)^23^. Key residues (orange) and lipid-binding residues (dark red) that exhibit deficits in cellular necroptotic signaling are shown as sticks. The lower panel shows a representation of the mouse MLKL α1-α2 helix and α3-α4 helix molecular surface, where each residue is color-coded as above. **c)** Cartoon representation of human MLKL 4HB (grey) + first brace (dark grey, Top panel) (PDB, 2MSV)^43^. Key IP6-binding residues (purple) and lipid-binding residues (dark red) that exhibit deficits in cellular necroptotic signaling are shown as sticks. The lower panel shows a representation of the human MLKL α1-α2 helix and α3-α4 helix molecular surface, where each residue is color-coded as above. Between the surface representations in (**b**) and (**c**), the different lipid-binding epitope for mouse and human MLKL can be observed.

By mapping the crucial residues for necroptotic signaling identified herein, and from earlier studies^28, 37^, on to the mouse MLKL 4HB domain+brace structure, it emerges that the lipid-binding epitope is centered on the α3-α4 helical face of mouse MLKL (**Figure 5b**). Importantly, this differs from the epitope in human MLKL, which is located on the opposing α1-α2 helical face of the 4HB domain (**Figure 5a, c**), as deduced from a combination of NMR spectroscopy, liposome permeabilization assays and cell death assays^29, 42, 43, 47^. In contrast to mouse MLKL, the implicated residues in human MLKL lipid interaction and membrane permeabilization are typically positively-charged (**Figure 5a, c; Supplementary Table 2**). While no studies have been performed on the capacity of mouse MLKL 4HB domain to engage inositol phosphates to date, it is notable that, again, positively charged residues within the human MLKL 4HB domain have been attributed functions in inositol phosphate binding. Although the sites of lipid engagement employed by the 4HB domain of plant MLKL are currently unknown^52^, it is notable that basic residues are largely absent from regions corresponding to binding residues within human MLKL. In our study of mouse MLKL, we used a plasma membrane-like lipid cocktail (**Supplementary Table 4**) to prepare unilamellar vesicles termed liposomes for our NMR titrations. On the other hand, studies of human MLKL have typically used isolated, highly negatively-charged phospholipid headgroups or inositol phosphates^40, 41, 42^, which may favour binding to positive sites on the human MLKL 4HB domain. Additionally, the inositol phosphate binding epitope overlaps that of the lipid-binding epitope and, as a result, this poses challenges for ascribing clear functions for inositol phosphates in regulating necroptotic signaling in cells. The precise function of inositol phosphates as modulators of necroptosis, and whether a similar regulatory function is conferred upon mouse MLKL, remains of outstanding interest.

Collectively, our findings uncover distinct species-dependent differences in lipid recognition between mouse and human MLKL. This plasticity illustrates the role of the HeLo/4HB domain as a scaffold for lipid engagement and permeabilization. Importantly, this work provides invaluable insight into how MLKL mediates necroptotic cell death and establishes a platform for future high-resolution structural studies in membranes to address the precise mechanism by which MLKL permeabilizes membranes.

## ACKNOWLEDGEMENTS

We thank the NMR facility (University of Melbourne), which is enabled by an Australian Research Council equipment grant LE120100022. We are grateful to the National Health and Medical Research Council for fellowship (J.M.H., 1142669; P.E.C., 1079700; J.M.M., 1105754, 1172929), grant (1057905; 1124735, 2002965) and infrastructure (IRIISS 9000653) support; and the Victorian Government Operational Infrastructure Support scheme. We acknowledge Australian Government Research Training Program Stipend Scholarships support (to SEG and AVJ) and the Wendy Dowsett Scholarship (to SEG). The National Deuteration Facility is partly funded by the National Collaborative Research Infrastructure Strategy (NCRIS), an Australian Government initiative.

## AUTHOR CONTRIBUTIONS

AS and CRH designed and performed experiments, analysed data and co-wrote the paper with JMM; KW carried out the biodeuteration of the recombinant protein for NMR; CF, KAD, SEG, AVJ, ALS, JMH and AW performed experiments and analysed data; PEC, EJP, PRG and JMM supervised the project and contributed to experimental design and data analysis. All authors commented on the manuscript.

## METHODS

### Expression constructs

For expression in mammalian cells, wild-type full-length MLKL was amplified by PCR from a mouse MLKL template (synthesised by DNA2.0, CA) and ligated into the doxycycline-inducible, puromycin-selectable mammalian expression vector, pF TRE3G PGK puro (Amp^r^) using BamHI and EcoRI restriction sites, as before^23^. Mutant mouse MLKL cDNAs were synthesized and subcloned into pF TRE3G PGK puro as BamHI-EcoRI fragments by ATUM (CA). Vector DNA was co-transfected into HEK293T cells with pVSVg and pCMV ΔR8.2 helper plasmids to generate lentiviral particles, which were transduced into three biologically independent Mouse Dermal Fibroblast (MDF) cell lines (*Mlkl*^*-/-*^, derived from different mice using a previously described method^23^) and selected for genomic integration using puromycin (2.5 μg mL^-1^; StemCell Technologies) using established procedures^28, 36^. For recombinant protein constructs, wild-type mouse MLKL_(1-158)_ was amplified by PCR from the mouse MLKL template and subcloned into the bacterial expression vector pETNusH Htb (Kan^r^) (derived from pETM60) ^53, 54^ as an in-frame fusion with a TEV (tobacco etch virus) protease-cleavable NusA-His_6_ tag. Mutant mouse MLKL_(1-158)_ constructs were amplified by PCR from the respective pF TRE3G PGK puro constructs (ATUM, CA) and subcloned into pETNusH Htb as BamHI-EcoRI fragments. All insert sequences were verified by Sanger sequencing (AGRF, VIC, Australia). All primers used in this study are listed in **Supplementary Table 3**.

### Biodeuteration and protein expression of mouse MLKL_(1-158)_

Uniformly ^15^N-labelled ([U-^15^N]), ^13^C^15^N-labelled [U-^13^C,^15^N] and fractional deuterated (f-^2^H) [U-^13^C, ^15^N]-labelled recombinant mouse MLKL_(1-158)_ was expressed via the pETNusH Htb vector at the National Deuteration Facility (NDF), Australian Nuclear Science and Technology Organization (ANSTO) in 1 L batch cultures using an established high cell density protocol^55^.

Briefly, for the [U-^15^N]- and [U-^13^C,^15^N]-labelled mouse MLKL_(1-158)_ constructs, 300 µL of freshly transformed *E. coli* BL21Star™(DE3) cells were inoculated into 10 mL of H_2_O ModC1 minimal medium, supplemented with kanamycin (40 µg L^-1^) and incubated overnight at 30 °C shaking at 220 rpm. The cell suspensions were diluted 5x in ^1^H, ^15^N-ModC1 medium (40 g/L glycerol, 5.16 g/L ^15^NH_4_Cl ≥ 98 atom % ^15^N) or ^1^H, ^13^C, ^15^N-ModC1 medium (20 g/L glycerol-^13^C_3_ 99 atom % ^13^C, 5.16 g/L ^15^NH_4_Cl ≥ 98 atom % ^15^N) and grown at 37 °C for two OD_600_ doublings, respectively for the [U-^15^N]) and [U-^13^C,^15^N] labelled constructs. Finally, cells were inoculated into fresh ^1^H, ^15^N-ModC1 to a volume of 100 mL and grown to an OD_600_ of 0.9-1.2 before inoculation into 900 mL of labelled expression medium (as described above) in a 1 L working volume bioreactor. *E. coli* cells were grown at 25 °C until OD_600_ of ∼9.5 and expression was induced by addition of isopropylthio-β-D-galactopyranoside (IPTG) at a final concentration of 1 mM. After ∼24 h induction at 20 °C, during which a further 5.16 g of ^15^NH_4_Cl was added to the culture, each labelled cell suspension was pelleted by centrifugation at 8000×*g* for 20 min and each biomass stored at −80 °C.

Briefly, a three step deuterated minimal medium adaptation process was followed for the (f-^2^H) [U-^13^C, ^15^N]-labelled mouse MLKL_(1-158)_ construct starting with 300 µL of freshly transformed *E. coli* BL21Star™(DE3) cells inoculated into 10 mL of 50% deuterium oxide (D_2_O) (v/v) ModC1 minimal medium (20 g/L glycerol) with 40 µg L^-1^ kanamycin and incubated overnight at 37 °C with shaking at 220 rpm. The resulting cell suspension was diluted 5-fold in ^2^H, ^13^C, ^15^N-ModC1 medium (D_2_O 99.8 atom % D, 20 g/L glycerol-^13^C_3_ 99 atom % ^13^C, 5.16 g/L ^15^NH_4_Cl ≥ 98 atom % ^15^N) and grown at 37 °C for approximately one OD_600_ doubling. Finally, cells were inoculated into fresh ^2^H, ^13^C, ^15^N-ModC1 to a volume of 100 mL and grown to an OD_600_ of 1.1 before inoculation into 900 mL of labelled expression medium as described in a 1 L working volume bioreactor. *E. coli* cells were grown at 37 °C until OD_600_ reached 8.2 and expression induced by addition of IPTG at a final concentration of 1 mM. After 26 h induction at 20 °C, during which a further 5.16 g of ^15^NH_4_Cl was added to the culture, the labelled cell suspension was pelleted by centrifugation at 8000×*g* for 20 min and biomass stored at −80 °C.

### Protein expression of unlabelled mouse MLKL (1-158)

Mouse MLKL_(1-158)_ constructs with in-frame TEV protease cleavable N-terminal NusA-His_6_ tags (pETNusH Htb) were expressed in *E. coli* BL21-Codon Plus (DE3)-RIL cells cultured in Super Broth supplemented with kanamycin (50 µg mL^-1^) at 37°C with shaking at 220 rpm to an OD_600_ of ∼0.6-0.8. Protein expression was induced by the addition of IPTG (final concentration of 1 mM) and the temperature was lowered to 18 °C for incubation overnight. Following protein expression, the cell suspension was pelleted by centrifugation at 8000×*g* for 20 min and biomass stored at −80 °C.

### Recombinant protein purification

For liposome permeabilization assays, cell pellets of mouse MLKL_(1-158)_ were resuspended in wash buffer [20 mM Tris-HCl (pH 8.0), 500 mM NaCl, 5 mM imidazole (pH 8.0), 20% glycerol, 1 mM TCEP [Tris-(2-carboxyethyl)phosphine], supplemented with Complete protease inhibitor cocktail (Roche), and lysed by sonication. The whole cell lysate was clarified by centrifugation (45,000×*g*, 1 h, 4 °C), filtered (0.2 μM) and the supernatant was incubated with pre-equilibrated Ni-NTA agarose (HisTag, Roche) at 4 °C for 1 h with gentle agitation. Ni-NTA beads were then pelleted via centrifugation and washed thoroughly with wash buffer. Bound protein was eluted from the beads using elution buffer [20 mM Tris-HCl (pH 8.0), 500 mM NaCl, 250 mM imidazole (pH 8.0), 20% glycerol, 1 mM TCEP], filtered through a 0.45-μm filter, mixed with 300 μg of recombinant His_6_-TEV and dialysed overnight in size exclusion buffer [20 mM Tris-HCl (pH 8.0), 150 mM NaCl, 1 mM TCEP], supplemented with 5% glycerol at 4 °C. Following protease cleavage, the dialysate was further purified using Ni-NTA chromatography to eliminate uncut material, cleaved NusA and the TEV protease. The flowthrough containing the mouse MLKL_(1-158)_ construct was concentrated via centrifugal ultrafiltration (10 kDa molecular weight cut-off; Millipore) and loaded onto a Superdex 75 Increase 10/300 GL size exclusion column (Cytiva) equilibrated in size exclusion buffer. Protein purity was assessed by SDS-PAGE (**Fig. 3c**). Protein that was not immediately used in experiments was aliquoted, flash-frozen in liquid nitrogen and stored at −80 °C.

For NMR experiments, cell pellets of [U-^15^N]-, [U-^13^C,^15^N]- and (f-^2^H) [U-^13^C, ^15^N]-labelled mouse MLKL_(1-158)_ were resuspended in low imidazole buffer [20 mM HEPES (pH 7.5), 200 mM NaCl, 5% v/v glycerol, 35 mM imidazole (pH 7.5), 1 mM TCEP], supplemented with Complete protease cocktail inhibitor (Roche), and lysed by sonication. The whole cell lysate was clarified by centrifugation (45,000×*g*, 1 h, 4 °C), filtered (0.2 μM) and the supernatant was loaded onto a HisTrap FF 5 ml column (Cytiva) pre-equilibrated with low imidazole buffer at 4 °C. After washing in low imidazole buffer, the bound protein was eluted using high imidazole buffer [20 mM HEPES (pH 7.5), 200 mM NaCl, 5% v/v glycerol, 375 mM imidazole (pH 7.5), 1 mM TCEP]. The eluant was further purified by cleaving the NusA-His_6_ tag by incubating with TEV protease, dialysis overnight in size exclusion buffer [20 mM HEPES (pH 7.5), 200 mM NaCl, 1 mM TCEP] and a second round of HisTrap-chromatography to eliminate uncut material, cleaved NusA and the TEV protease. The flowthrough was concentrated via centrifugal ultrafiltration (10 kDa molecular weight cut-off; Millipore) and loaded onto a Superdex 75 Increase 10/300 GL size exclusion column (Cytiva) equilibrated in NMR size exclusion buffer [20 mM HEPES (pH 6.8), 100 mM NaCl, 1 mM TCEP]. Protein purity was assessed by SDS-PAGE and then used fresh for each NMR experiment.

### NMR samples and NMR Spectroscopy

NMR experiments were all performed at 25 °C on a 700-MHz Bruker Avance HDIII spectrometer equipped with triple resonance cryoprobe. Proteins samples were prepared in a buffer containing 20 mM HEPES, 100 mM NaCl and 1 mM TCEP at pH 6.8 supplemented with 10% ^2^H_2_O. Backbone resonances (^13^Cα, ^13^Cβ, ^13^C’
s, ^15^N and NH) of residues were assigned from 3D HNCACB, HN(CO)CACB, HNCO and HNCA experiments using non-uniform sampling (NUS). For NUS, sampling schedules were generated using Poisson gap sampler with 10% of the total number of points collected for all the 3D NMR experiments^56^. Spectra were reconstructed with compressed sensing algorithm using qMDD^57^ and processed using NMRPipe^58^ and data analyzed in NMRFAM-SPARKY^59^. The ^1^H chemical shifts were referenced directly to DSS at 0 ppm and the ^13^C and ^15^N chemical shifts were subsequently referenced using the ^13^C/^1^H and ^15^N/^1^H ratios as described previously^60^.

### NMR relaxation experiments

Protein was used at ∼180 µM with 2.5 mM liposomes (100 nm diameter). TCEP was added fresh (to 1mM) to sample prior to data collection. NMR size exclusion buffer [20 mM HEPES (pH 6.8), 100 mM NaCl, 1 mM TCEP]. ^15^N-R_2_ experiments were collected with a recycle time of 2.6 s and 16 scans per FID and ^15^N{^1^H}-NOE experiments were collected with a saturation pulse of 4 s and an additional relaxation delay of 5 s and 32 scans per FID. ^15^N-R_2_ relaxation delays of 16.96 (×2), 33.92, 67.84, 101.76 (×2), 135.68, 169.6 (×2), 203.52 and 237.44 ms were used. The repeated spectra were used to estimate instrumental error. ^15^N relaxation parameters were determined using the program *relax* (version 3.3.4)^61^. For R_2_ rate constants, errors were estimated using 500 Monte Carlo Simulations. The steady state ^15^N{^1^H}-NOE values for mouse MLKL_(1-158)_ were estimated from the ratios of peak intensities obtained from spectra acquired with and without proton saturation using *relax*. Errors for ^15^N{^1^H}-NOE experiment were calculated based on noise level in the spectrum.

### Reagents and antibodies

Primary antibodies used in this study for immunoblotting were: rat anti-mouse MLKL (WEHI clone 5A6; produced in-house and soon available from Millipore as MABC1634; 1:2000)^49^; rabbit anti-phospho-S345 mouse MLKL (Cell Signaling Technology; clone D6E3G; 1:2000); rat anti-mouse RIPK3 (WEHI clone 8G7; produced in-house^7^ and soon available from Millipore as MABC1595; 1:2000); rabbit anti-phospho-T231/S232 mouse RIPK3 (Genentech; clone GEN135-35-9 ^62^; lot PUR73907; 1:2000); rabbit anti-mouse or human RIPK1 (Cell Signaling Technology; clone D94C12; 1:2000); and mouse anti-Actin (A1978, Sigma-Aldrich, St Louis, MO, USA; 1:5000). Secondary antibodies used in this study were: horseradish peroxidase (HRP)-conjugated goat anti-rat IgG (Southern Biotech 3010-05), HRP-conjugated goat anti-mouse IgG (Southern Biotech 1010-05), and HRP-conjugated goat anti-rabbit IgG (Southern Biotech 4010-05). All secondary antibodies were used at a dilution of 1:10000. Recombinant hTNF-Fc, produced in-house, and the Smac-mimetic, Compound A, have been previously described^63, 64^. The pan-caspase inhibitor, IDN-6556/emricasan, was provided by Tetralogic Pharmaceuticals.

### Cell culture

*Mlkl*^*-/-*^ MDF cells were cultured in Dulbecco’s Modified Eagle Medium (DMEM; Gibco) supplemented with 8% (v/v) Fetal Calf Serum (FCS; Sigma), penicillin (100 U mL^-1^), streptomycin (100 μg mL^-1^). Puromycin (2.5 μg mL^-1^; StemCell Technologies) was added for lines stably transduced with inducible mouse MLKL constructs. Routine testing confirmed cell lines to be mycoplasma-negative.

### IncuCyte cell death assays

*Mlkl*^*-/-*^ MDF cells were seeded into 96-well plates at 8 ×10^4^ cells/well and left to adhere for 4-5 h prior to treatment with doxycycline (20 ng mL^-1^) overnight to induce expression of the relevant full-length mouse MLKL constructs. Cells were then treated with necroptotic stimulus comprising, TNF (100 ng mL^-1^), the Smac-mimetic compound A (500 nM) and the pan-caspase inhibitor IDN-6556 (10 μM) (TSI) to induce necroptosis in FluoroBrite DMEM media (ThermoFisher Scientific) supplemented with 1% FCS, 1 mM Na pyruvate (ThermoFisher Scientific), 1 mM L-GlutaMAX (ThermoFisher Scientific), SYTOX Green nucleic acid stain (ThermoFisher Scientific, 1:20000) and SPY620 live cell DNA stain (Spirochrome, 1:1000). Cells were then imaged using the IncuCyte S3 System (Essen Bioscience) with default bright-field, red and green channel settings on 10x objective. Scans were obtained every 30 min for 5 h, where percent cell death was quantified based upon the number of SYTOX Green-positive cells per image over the number of SPY620-positive cells per image using IncuCyte S3 v2018A software (Essen Bioscience). Data were plotted as mean ± SEM from three biologically independent *Mlkl*^*-/-*^ MDF cell lines (*n* = 6 to 9).

### Immunoblot

*Mlkl*^*-/-*^ MDF cells were seeded into 24-well plates at 7 ×10^4^ cells/well and left to adhere for 4-5 h. Cells were induced overnight with doxycycline (20 ng mL^-1^) and then treated with TNF (100 ng mL^-1^), Smac-mimetic (Compound A; 500 nM) and pan-caspase inhibitor, IDN-6556 (5 μM) (TSI) for 1.5 or 3.0 h. Cells were lysed in ice-cold RIPA buffer [10 mM Tris-HCl pH 8.0, 1 mM EGTA, 2 mM MgCl_2_, 0.5% v/v Triton X100, 0.1% w/v Na deoxycholate, 0.5% w/v SDS and 90 mM NaCl] supplemented with 1x Protease & Phosphatase Inhibitor Cocktail (Roche) and 100 U/mL Denarase (c-LEcta). Whole-cell lysates were boiled at 100 °C for 10-15 min in 1 × SDS Laemmli lysis buffer (126 mM Tris-HCl, pH 8, 20% v/v glycerol, 4% w/v SDS, 0.02% w/v bromophenol blue, 5% v/v 2-mercaptoethanol), and then resolved by 4 to 15% Tris-Glycine gel (Bio-Rad). Proteins were transferred to PVDF membrane, blocked with 5% w/v skim milk powder in TBST and then probed overnight with primary antibodies (as per *Reagents and antibodies* above). The signals were revealed by enhanced chemiluminescence on a ChemiDoc Touch Imaging System (BioRad) using an appropriate HRP-conjugated secondary antibody (as per *Reagents and antibodies* above). Before probing different proteins with primary antibody, membranes were incubated in mild stripping buffer [200 mM glycine pH 2.9, 1% w/v SDS, 0.5 mM TCEP] for 30 min at room temperature then re-blocked.

### Lactate Dehydrogenase (LDH) release

Colorimetric LDH release assay kit (Promega G1780) was performed according to manufacturer’s instructions. Data are plotted as mean ± SD of three independent replicates.

### Liposome preparation

Large Unilamellar Vesicles (LUVs) were prepared using a plasma membrane-like lipid mix (**Supplementary Table 4**) and resuspended in chloroform as a 20 mg mL^-1^ (∼25 mM for most lipids) stock as previously reported ^29, 37^. Dried lipids were resuspended in either 500 μL of LUV buffer [10 mM HEPES pH 7.5, 135 mM KCl] with 50 mM 5(6)-Carboxyfluorescein dye (Sigma) to form dye filled liposomes for permeabilization assays or NMR size exclusion buffer [20 mM HEPES (pH 6.8), 100 mM NaCl, 1 mM TCEP] for NMR studies. The mixture was then freeze-thawed at least 5×, by immersion in liquid nitrogen until fully frozen, followed by immersion in a 37 °C water bath until contents had thawed. The lipid mixture was extruded through polycarbonate membranes of 100 nm size cut-off (Avanti Polar Lipids, AL, USA), a minimum of 21 times to form liposomes, using a pre-warmed mini extruder (Avanti Polar Lipids, AL, USA). Liposomes stocks were at approximately 2.5 mM lipid concentration and were stored at 4 °C in the dark.

### Liposome dye release assays

Recombinant mouse MLKL_(1-158)_ protein was diluted to 16 μM (2× desired final concentration) in LUV buffer, and 50 μL aliquoted into adjacent wells of a 96 well Flat-bottom plate (ThermoFisher Scientific). Prior to use, the liposomes (100 nm diameter filled with 5(6)-Carboxyfluorescein dye) were purified from excess dye using a PD-10 desalting column (Cytiva) and diluted to 20 μM in LUV buffer. At the plate reader (Hidex Chameleon Multilabel Microplate Reader; Lab Logic), the protocol was pre-programmed, before 50 μL of liposomes was promptly added to each well of the 96 well plate using a multi-channel pipette. The plate was then immediately placed in the plate reader and measurements started. Fluorescence was measured every 2 min for 60 minutes (31 measurements) at 20 °C with excitation wavelength of 485 nm and emission wavelength of 535 nm. 100% dye release was determined by the incubation of liposomes with 50 μL of 1% CHAPS detergent in LUV buffer, while a baseline was determined by the incubation of liposomes with 50 μL of LUV buffer alone. All assays were performed in triplicate. Data were plotted as mean ± SEM of three independent assays and data is presented as a percentage of maximum dye release.

