## Supplementary Information for "Membrane permeabilization is mediated by distinct epitopes in mouse and human orthologs of the necroptosis effector, MLKL"

### SUPPLEMENTARY FIGURES

**Supplementary Figure 1 | Expression of wild-type and mutant mouse MLKL in *Mlkl*$^{-/-}$ MDF cells.** Following doxycycline (Dox) induction, whole cell-lysates were fractioned by SDS-PAGE and probed by immunoblot for MLKL with anti-actin as a loading control. Immunoblots are representative of $n = 2$ independent experiments.

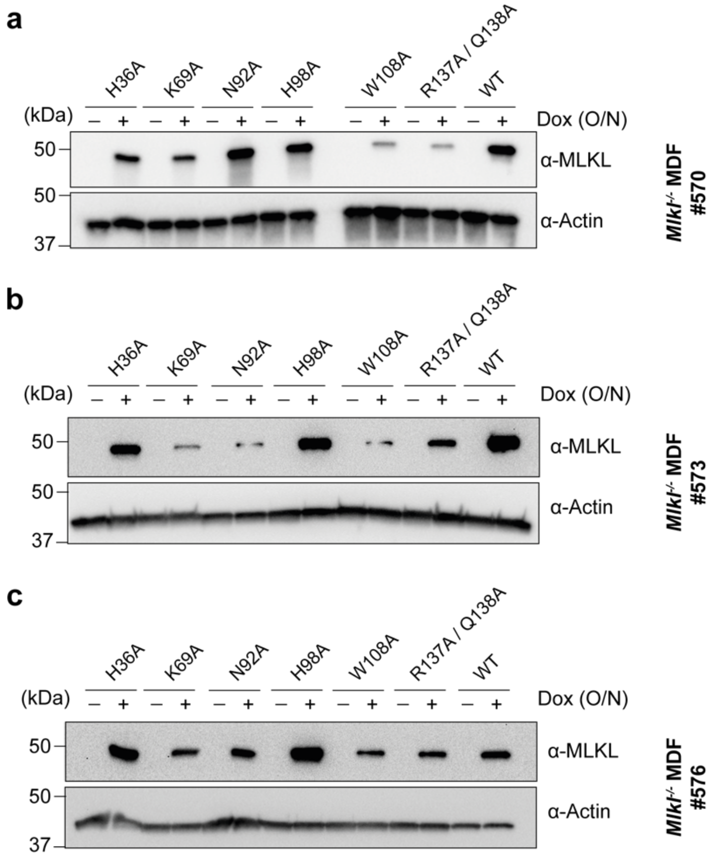

**Supplementary Figure 2 | Raw IncuCyte data of cell death mediated by wild-type and mutant mouse MLKL. a-f)** To establish the contribution of each lipid-binding residue in cellular necroptosis signaling, full-length wild-type (WT) and mutant mouse MLKL were stably introduced into *Mlkl*$^{-/-}$ MDF cells. Following doxycycline (Dox) treatment to induce expression, percent cell death was quantified using IncuCyte S3 live cell imaging in the presence or absence of the necroptotic stimulus, TNF and Smac-mimetic Compound A and pan-caspase inhibitor, IDN-6556, (TSI) for 5 h, by determining the number SYTOX Green-positive cells (dead cells) relative to the number of SPY620-positive cells (total cell confluency). The percent cell death of wild-type mouse MLKL is shown in each plot as a reference. Data represent mean ± SEM from three biologically independent *Mlkl*$^{-/-}$ MDF cell lines ($n$ = 6 to 9).

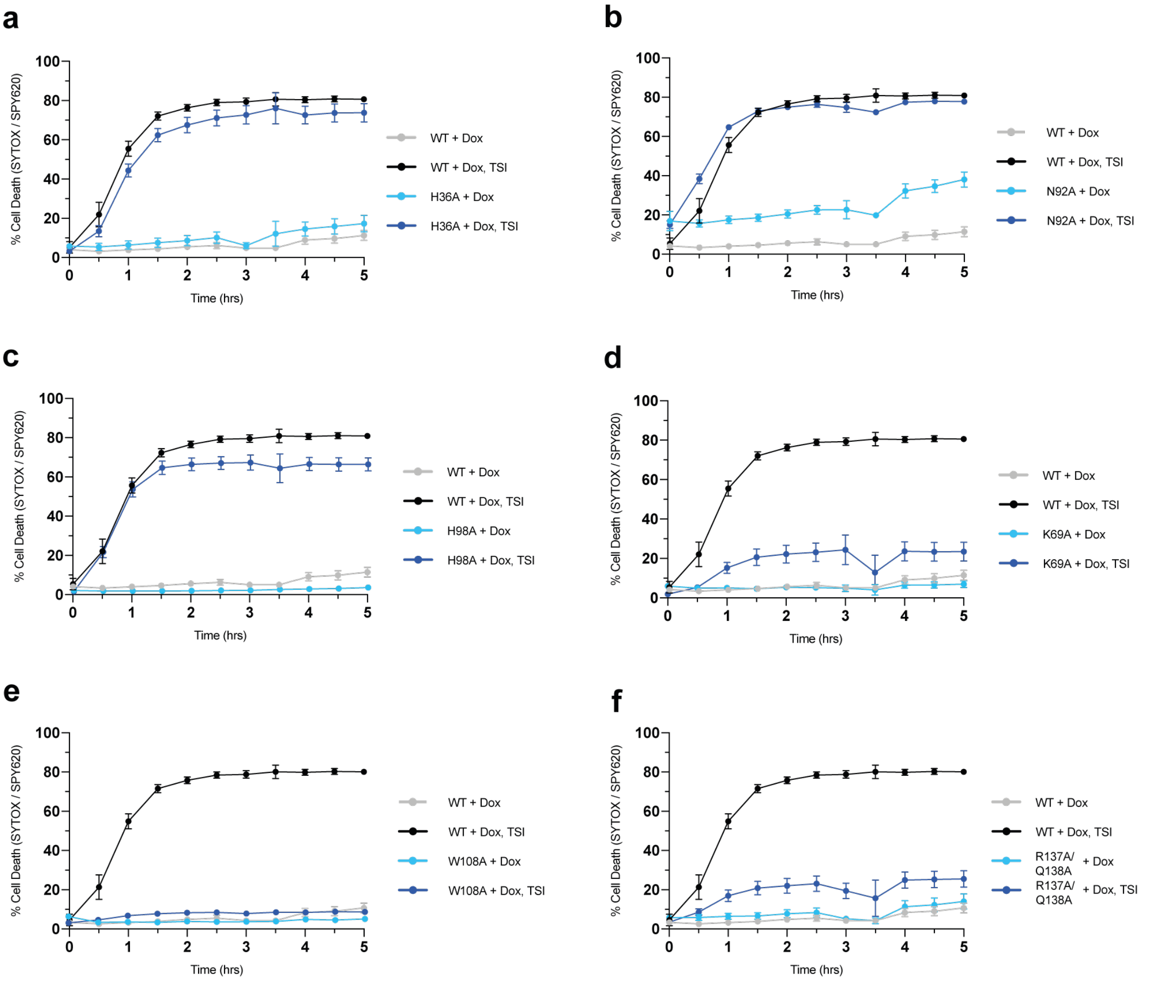

### SUPPLEMENTARY TABLES

#### Supplementary Table 1 | Summary of wild-type and mutant mouse MLKL properties

| MLKL mutation | Location of mutation | Lipid-binding (Impact on liposome permeabilization) | Necroptotic signalling function |
|---|---|---|---|
| mMLKL WT | | | + |
| mMLKL Y15A/E16A | α1 helix | N/D | -[1] |
| mMLKL C18A/C24A/C28A | α1/2 helix | N/D | -[1] |
| mMLKL K22A/R30A | α2 helix and preceding loop | N/D | -[1] |
| mMLKL H36A | α2 helix | Yes (compromised) | +[†] |
| mMLKL R63A/D65A | α3 helix | N/D | -[1] |
| mMLKL K69A | α3 helix | Yes (compromised) | Reduced[†] |
| mMLKL E70A/N72A | α3 helix | N/D | -[1] |
| mMLKL E76A/K77A | α3 helix | N/D | -[1] |
| mMLKL K80A/K81A | α3-α4 loop | N/D | -[1] |
| mMLKL N92A | α3-α4 loop | Yes (comparable to WT) | +[†] |
| mMLKL H98A | α4 helix | Yes (compromised) | +[†] |
| mMLKL H98A/E99A | α4 helix | N/D | -[1] |
| mMLKL E102A/K103A | α4 helix | N/D | -[1] |
| mMLKL R105A/D106A | α4 helix | N/D | -[1, 2] |
| mMLKL W108A | α4 helix | Yes (compromised) | -[†] |
| mMLKL E109A/E110A | α4 helix | N/D | -[1, 2] |
| mMLKL LLLL$^{112-115}$AAAA | α4 helix | N/D | -[1] |
| mMLKL R137A/Q138A | First brace helix | Yes (compromised) | -[†] |

N/D = Not determined; - = loss-of-function; [†] This study

**Supplementary Table 2 | Summary of wild-type and mutant human MLKL properties**

| MLKL mutation | Location of mutation | Lipid- or IP6-interactor | Impact on liposome permeabilization | Necroptotic signalling function |
|---|---|---|---|---|
| hMLKL WT | | | | + |
| hMLKL E2A/N3A | α1 helix | Lipid | Compromised | -[3] |
| hMLKL K5A | α1 helix | Lipid | Compromised | -[3] |
| hMLKL H15A | α1 helix | IP6 | N/D | N/D[4] |
| hMLKL K16A/R17A | α1 helix | Lipid | Compromised | Reduced[3, 5] |
| hMLKL E19A | α1 helix | IP6 | N/D | N/D[6] |
| hMLKL K22Q/K25Q | α2 helix and preceding loop | Lipid | Compromised | -[7] |
| hMLKL R29E/R30E | α2 helix | Lipid | Compromised | -[7] |
| hMLKL L36A | α2 helix | IP6 | N/D | N/D[6] |
| hMLKL K50A/K51A | α2-α3 loop | Lipid | Compromised | -[3] |
| hMLKL K78A | α3 helix | IP6 | N/D | N/D[6] |
| hMLKL D107A/E111A | α4 helix | Lipid | Compromised | -[5, 8] |
| hMLKL L114A | α4 helix | Lipid | Compromised | -[8] |
| hMLKL L116A | α4 helix | IP6 | N/D | N/D[6] |
| hMLKL E119A | α4 helix | IP6 | N/D | N/D[6] |
| hMLKL R152A | First brace helix | IP6 | N/D | N/D[6] |

N/D = Not determined; - = loss-of-function

**Supplementary Table 3 | Primers used in this study**

| Primer name | Sequence |
|---|---|
| mMLKL Bam 5´ fwd* | 5´-CGCGGATCCatggataaattgggacagatcatc-3´ |
| mMLKL 158 stop EcoRI rev | 5´-CGCGAATTCAgctaatttgcaactgcatcaggataac-3´ |
| mMLKL 464 stop EcoRI rev | 5´-CGGAATTCttacaccttcttgtccgtggattc-3´ |

*Restriction sites underlined

**Supplementary Table 4 | Plasma membrane-like lipid mix for liposomes**

| Lipid | Proportion of plasma membrane-like mix | Source |
|---|---|---|
| Phosphatidylethanolamine (POPE) | 20% | Avanti Polar Lipids (Alabaster, AL, USA) |
| Phosphatidylcholine (POPC) | 40% | |
| Phosphatidylinositol (PI) | 10% | |
| Phosphatidylserine (DOPS) | 20% | |
| Phosphatidylglycerol (POPG) | 10% | |
